# Parallel phosphoproteomics and metabolomics map the global metabolic tyrosine phosphoproteome

**DOI:** 10.1101/2024.05.14.594136

**Authors:** Alissandra L. Hillis, Tigist Tamir, Grace E. Perry, John M. Asara, Jared L. Johnson, Tomer M. Yaron, Lewis C. Cantley, Forest M. White, Alex Toker

## Abstract

Tyrosine phosphorylation of metabolic enzymes is an evolutionarily conserved post-translational modification that facilitates rapid and reversible modulation of enzyme activity, localization or function. Despite the high abundance of tyrosine phosphorylation events detected on metabolic enzymes in high-throughput mass spectrometry-based studies, functional characterization of tyrosine phosphorylation sites has been limited to a subset of enzymes. Since tyrosine phosphorylation is dysregulated across human diseases, including cancer, understanding the consequences of metabolic enzyme tyrosine phosphorylation events is critical for informing disease biology and therapeutic interventions. To globally identify metabolic enzyme tyrosine phosphorylation events and simultaneously assign functional significance to these sites, we performed parallel phosphoproteomics and polar metabolomics in non-tumorigenic mammary epithelial cells (MCF10A) stimulated with epidermal growth factor (EGF) in the absence or presence of the epidermal growth factor receptor (EGFR) inhibitor erlotinib. We performed an integrated analysis of the phosphoproteomic and metabolomic datasets to identify tyrosine phosphorylation sites on metabolic enzymes with functional consequences. We identified two previously characterized (PKM, PGAM1) and two novel (GSTP1, GLUD1) tyrosine phosphorylation sites on metabolic enzymes with purported functions based on metabolomic analyses. We validated these hits using a doxycycline-inducible CRISPR interference (CRISPRi) system in MCF10A cells, in which target metabolic enzymes were depleted with simultaneous re-expression of wild-type, phosphomutant or phosphomimetic isoforms. Together, these data provide a framework for identification, prioritization and characterization of tyrosine phosphorylation sites on metabolic enzymes with functional significance.

## Introduction

Protein phosphorylation is an energy-efficient and reversible post-translational modification that regulates nearly every cellular process, including protein activity, localization and interactions^1–4^. Phosphorylation is catalyzed by protein kinases, and there are approximately 550 protein kinases in the human genome^1,4^. Given the low abundance of tyrosine phosphorylation (pTyr), there are only 90 tyrosine kinases in humans, including receptor (RTK) and non-receptor (NRTK) tyrosine kinases^5,6^. The epidermal growth factor receptor (EGFR) is an RTK that rapidly phosphorylates target proteins in response to epidermal growth factor (EGF)^6,7^. Protein tyrosine phosphatases (PTPs) catalyze the dephosphorylation of pTyr residues, and the coordinated action of tyrosine kinases and PTPs ensure proper pTyr signaling^5,8^.

Dysregulated protein phosphorylation is a hallmark of many diseases, including cancer. Despite its low abundance, pTyr has emerged as a key modification downstream of activated tyrosine kinases in cancer^2^. Tyrosine kinases are frequently mutated or overexpressed in cancer, resulting in the classification of at least 30 of the 90 tyrosine kinases as oncogenic^9^. Similarly, PTPs are often lost or repressed in cancer to sustain oncogenic pTyr events^5,8,10^. As a result, tyrosine kinases have become prominent drug targets, and many tyrosine kinase inhibitors (TKIs) have been approved for use in cancer treatment^11^. While the physiological importance of phosphorylation, including pTyr, is well recognized, our understanding of the molecular consequences of phosphorylation events in normal and pathophysiological states is limited^12^.

Among the proteins regulated by pTyr are metabolic enzymes. While phosphorylation sites in general are not evolutionarily conserved, phosphorylation sites on metabolic enzymes are highly positionally conserved^2,13^. This suggests that phosphorylation plays a key role in regulating the activity of metabolic enzymes. Tyrosine phosphorylation regulates the activity of metabolic enzymes by three known mechanisms. First, by introducing negative charges, phosphorylation can generate attractive or repulsive forces that can modify the positioning of amino acid side chains in the catalytic site, resulting in altered enzyme catalysis or altered affinity of the enzyme for its substrate^9^. Lactate dehydrogenase A (LDHA) and phosphoglycerate mutase 1 (PGAM1) are regulated by pTyr in this way^9,14–16^. Second, pTyr can affect the three-dimensional structure of metabolic enzymes by promoting the assembly or dissociation of enzyme subunits. pTyr of hexokinase 1/2 (HK1/2) and pyruvate kinase (PKM) regulates their multimerization^17–23^. Third, pTyr can modulate the intracellular localization of metabolic enzymes and their interactions with effector molecules. Phosphofructokinase 1 (PFK1) and glucose-6-phosphate dehydrogenase (G6PD) are regulated by pTyr in this way^9,24,25^.

While there are only a few notable examples of metabolic enzyme regulation by pTyr, global analyses indicate that there are hundreds of uncharacterized pTyr sites on metabolic enzymes that could regulate their activity. For example, a phosphoproteomics study in rat lungs identified 88 EGF-regulated pTyr sites, of which 12.5% were on metabolic proteins, suggesting that pTyr regulates metabolic enzymes *in vivo*^12^. Yet, little is known about how global changes in phosphorylation affect the regulation of metabolic enzymes and subsequent changes in cellular metabolism. The development of phosphoproteomics and pTyr enrichment methods has facilitated the identification of kinase-associated substrates, but phosphoproteomics alone cannot provide information about the functional consequences of protein phosphorylation. To simultaneously identify pTyr sites on metabolic enzymes and assign functional significance to these events, we performed parallel phosphoproteomics and polar metabolomics in non-tumorigenic mammary epithelial cells (MCF10A) stimulated with EGF in the absence or presence of the EGFR inhibitor erlotinib. An integrated analysis of the phosphoproteomic and metabolomic data identified functionally relevant, EGF-induced pTyr sites on metabolic enzymes.

## Results

### Phosphoproteomics and metabolomics map the altered metabolic tyrosine phosphoproteome in response to EGF stimulation

Tyrosine phosphorylation sites on metabolic enzymes are evolutionarily conserved, yet few of these sites have been functionally characterized. A search of the PhosphoSitePlus database for pTyr sites on metabolic enzymes with at least 1 low throughput study (LTP) identified pTyr sites on 78 unique metabolic enzymes^26^. Plotting these sites on a map of the human metabolome highlights that they are mostly limited to central carbon metabolism (Fig. 1*A*). However, expanding the PhosphoSitePlus search to include metabolic enzymes with pTyr sites detected in any study, including high throughput studies (HTPs), identified over 4000 pTyr sites across 1144 unique metabolic enzymes, involved in nearly every human metabolic process (Fig. 1*B*).^26^ These data show that only about 5% of the detected metabolic tyrosine phosphoproteome has been characterized.

**Figure 1.**
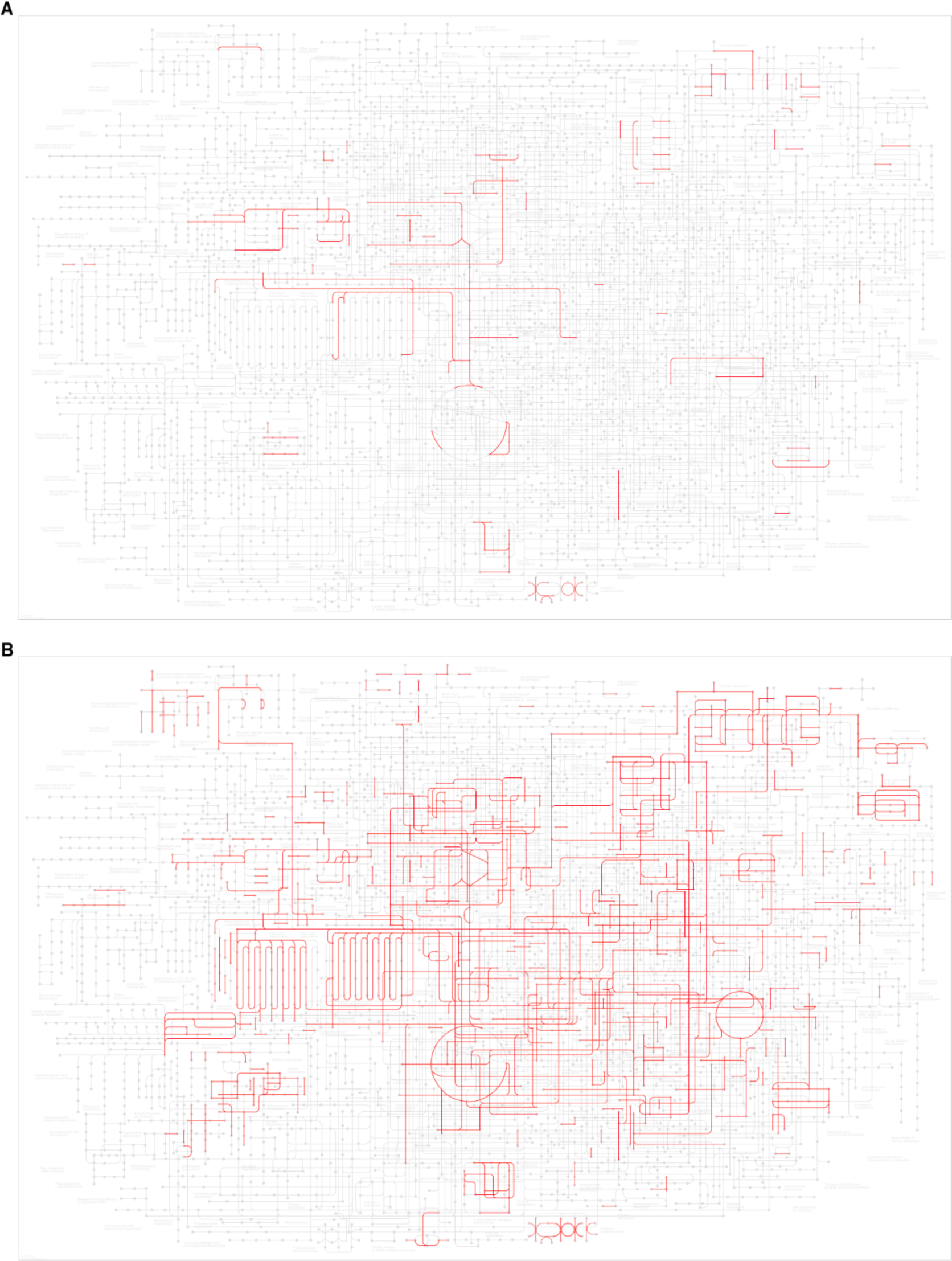
The metabolic tyrosine phosphoproteome is broadly uncharacterized. *A,* PhosphoSitePlus analysis of human metabolic enzymes with tyrosine phosphorylation (pTyr) sites that have been characterized by low throughput studies (LTPs). Enzymes with pTyr sites with greater than or equal to 1 LTP were plotted in red on a map of the human metabolome using KEGG Mapper (55 of 78 enzymes were mapped). *B,* PhosphoSitePlus analysis of human metabolic enzymes with pTyr sites that have been characterized by at least 1 high throughput study (HTP). Enzymes with pTyr sites with greater than or equal to 1 HTP were plotted in red on a map of the human metabolome using KEGG Mapper (562 of 1144 enzymes were mapped).

To characterize the metabolic tyrosine phosphoproteome in an unbiased manner, we performed parallel phosphoproteomics and metabolomics in non-tumorigenic mammary epithelial cells (MCF10A). MCF10A cells were stimulated with 10 ng/mL EGF for 5 minutes, 24 hours or 48 hours with or without a 10-minute pre-treatment with the EGFR inhibitor erlotinib or the PI3Kα inhibitor BYL719. We chose a 5-minute time point since EGFR signaling occurs rapidly after stimulation with EGF. However, metabolic enzymes may not be direct targets of EGFR, but rather targets of a NRTKs downstream of EGFR. We therefore also included 24- and 48-hour time points to capture phosphorylation events on metabolic enzymes that occur downstream of EGFR. Pre-treatment with BYL719 was utilized to inhibit the signaling contribution from PI3K driven activation of metabolism and enrich for EGF-mediated pTyr events downstream of EGFR. As expected, EGF stimulation increased the pTyr signature, which was potentiated with BYL719 pre-treatment and decreased with erlotinib pre-treatment (Fig. 2*A*). EGF stimulation resulted in phosphorylation of EGFR within 5 minutes. EGFR phosphorylation was rapidly attenuated, but phosphorylation of downstream targets such as AKT and ERK1/2 was sustained for 24 to 48 hours (Fig. 2*A*). Phosphorylation of EGFR and downstream targets was abolished with erlotinib pre-treatment and sustained with BYL719 pre-treatment (Fig. 2*A*). To verify that these time points were appropriate for identifying pTyr events on metabolic enzymes, we measured phosphorylation levels of well-characterized pTyr sites on PKM2 (Y105) and LDHA (Y10). Phosphorylation of PKM2 Y105 and LDHA Y10 was detected after EGF stimulation for 5 minutes, 24 hours or 48 hours (Fig. 2*A*). Together, these data support the relevance of these conditions for studying the functional consequences of pTyr events on metabolic enzymes.

**Figure 2.**
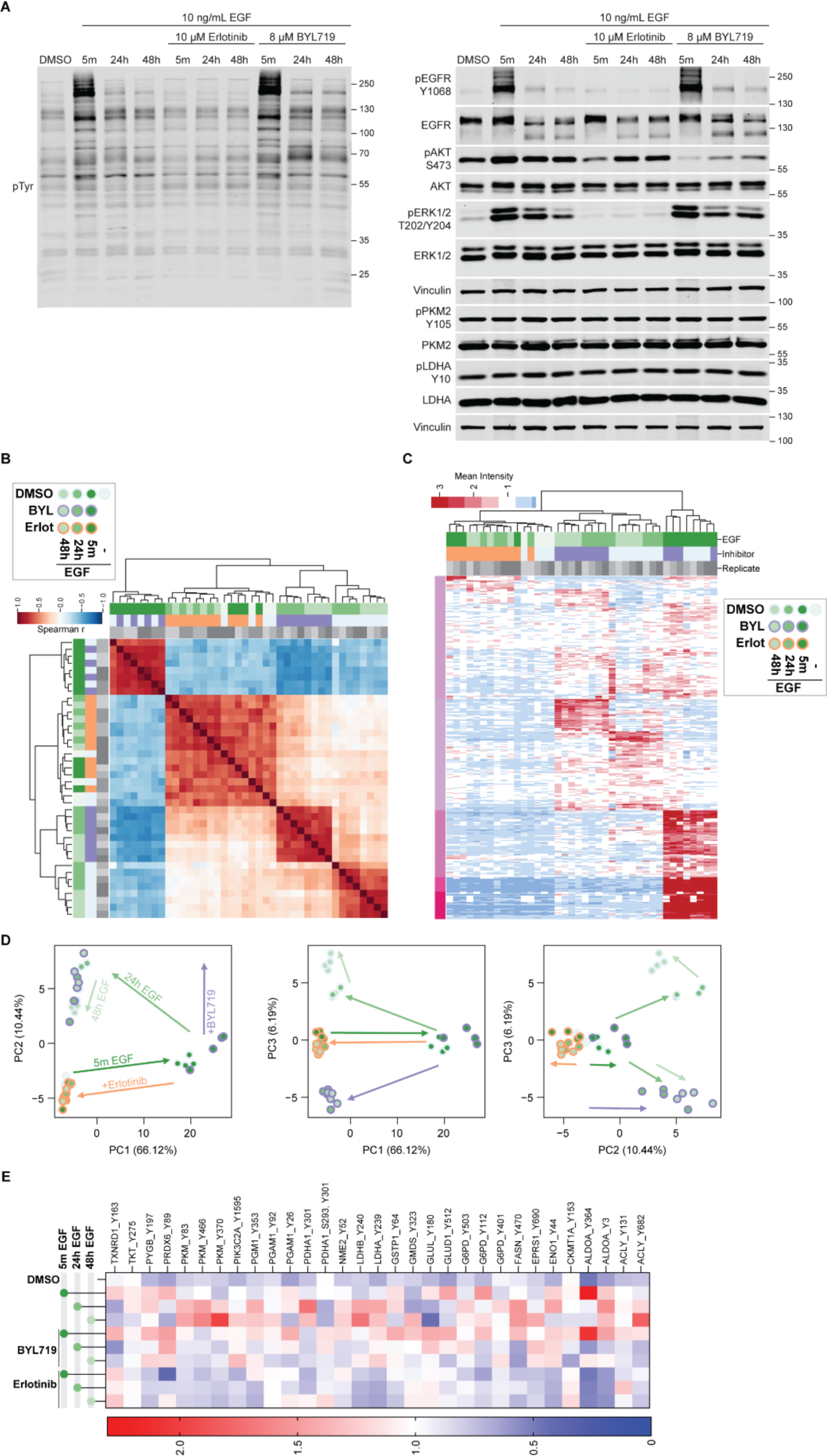
Phosphoproteomics in MCF10A cells identifies pTyr sites on metabolic enzymes downstream of activated EGFR. *A,* Immunoblots of total phosphotyrosine (pTyr), pEGFR^Y1068^, pAKT^S473^, pERK1/2^T202/Y204^, pPKM2^Y105^ and pLDHA^Y10^ in MCF10A cells after treatment with 10 ng/mL EGF for 5 minutes, 24 hours or 48 hours with or without a 10-minute pre-treatment with erlotinib (10 µM) or BYL719 (8 µM). Cells were seeded and harvested independently from the phosphoproteomics experiment in *B-C*. *B,* Hierarchical clustering based on the spearman correlation coefficients between phosphosites across all samples with a cutoff of +/-0.6 (distance metric = correlation) in MCF10A cells after treatment with 10 ng/mL EGF for 5 minutes, 24 hours or 48 hours with or without a 10-minute pre-treatment with erlotinib (10 µM) or BYL719 (8 µM). *C,* Heatmap showing fold-change in abundance of each phosphosite detected over the mean across all samples in MCF10A cells after treatment with 10 ng/mL EGF for 5 minutes, 24 hours or 48 hours with or without a 10-minute pre-treatment with erlotinib (10 µM) or BYL719 (8 µM). *D,* Principal Component Analysis (PCA) shows over 65% of the variance (PC1) in pTyr levels is driven by time-dependent EGF stimulation, while ∼10-15% of variance (PC2/PC3) is explained by BYL719 treatment. *E,* Heatmap showing fold-change in abundance of metabolic enzyme phosphosites detected over the mean across all samples in MCF10A cells after treatment with 10 ng/mL EGF for 5 minutes, 24 hours or 48 hours with or without a 10-minute pre-treatment with erlotinib (10 µM) or BYL719 (8 µM).

To quantitatively evaluate changes in pTyr on metabolic enzymes, we performed phosphoproteomics with pTyr enrichment in MCF10A cells stimulated with EGF for 5 minutes, 24 hours or 48 hours with or without a 10-minute pre-treatment with erlotinib or BYL719. Phosphoproteomic analysis identified strong correlation between samples stimulated with EGF for 5 minutes with or without BYL719-pre-treatment, suggesting that PI3Kα inhibition does not significantly alter the EGF-induced pTyr signature (Fig. 2*B-C*). Principal Component Analysis (PCA) revealed that the most significant changes in pTyr occurred within 5 minutes of EGF stimulation, as represented by PC1, which explains over 65% of the variance in pTyr levels (Fig. 2*D*). While most pTyr events were dampened 24-48 hours after EGF stimulation, a small proportion of phosphorylation events were sustained, as represented by PC2, which explains approximately 10% of the variance in pTyr levels (Fig. 2*D*). BYL719 pre-treatment promoted sustained pTyr phosphorylation out to 24 and 48 hours, represented by both PC2 and PC3, which likely reflects enhanced compensatory signaling through non-PI3K/AKT signaling pathways (Fig. 2*D*). We identified pTyr sites on 31 metabolic enzymes, with sites on 22 enzymes observed in at least 2 replicates (Fig. 2*E*). The majority of pTyr sites identified were on enzymes involved in glycolysis (Fig. 2*E*).

We next performed steady-state polar metabolomics in parallel with the phosphoproteomics. For metabolomics, we excluded the 5-minute time point since we did not expect to detect reliable changes in steady-state metabolite abundances at this short time point. Protein was harvested in parallel with metabolites for normalization. As in Fig. 2*A*, EGF stimulation for 24 and 48 hours induced global pTyr and phosphorylation of EGFR and its downstream targets AKT and ERK1/2, which was reversed with erlotinib pre-treatment (Fig. 3*A*). EGF stimulation also enhanced phosphorylation of metabolic enzymes (PKM2, LDHA), which was abrogated with erlotinib pre-treatment (Fig. 3*A*). For metabolomic analyses, we focused on changes in metabolite abundance with EGF stimulation compared to erlotinib pre-treatment. Although BYL719 pre-treatment enriched for pTyr events on metabolic enzymes, broad metabolic changes induced by PI3Kα inhibition would likely confound attempts to evaluate the functional consequences of metabolic enzyme pTyr sites. Metabolic pathways differentially regulated with EGF stimulation versus EGFR inhibition included glycolysis, nucleotide metabolism, glutathione metabolism, arginine metabolism and glutamine metabolism (Fig. 3*B-C*). To determine which metabolic changes could be attributed to altered pTyr of metabolic enzymes, we performed an integrated analysis of the phosphoproteomic and metabolomic datasets.

**Figure 3.**
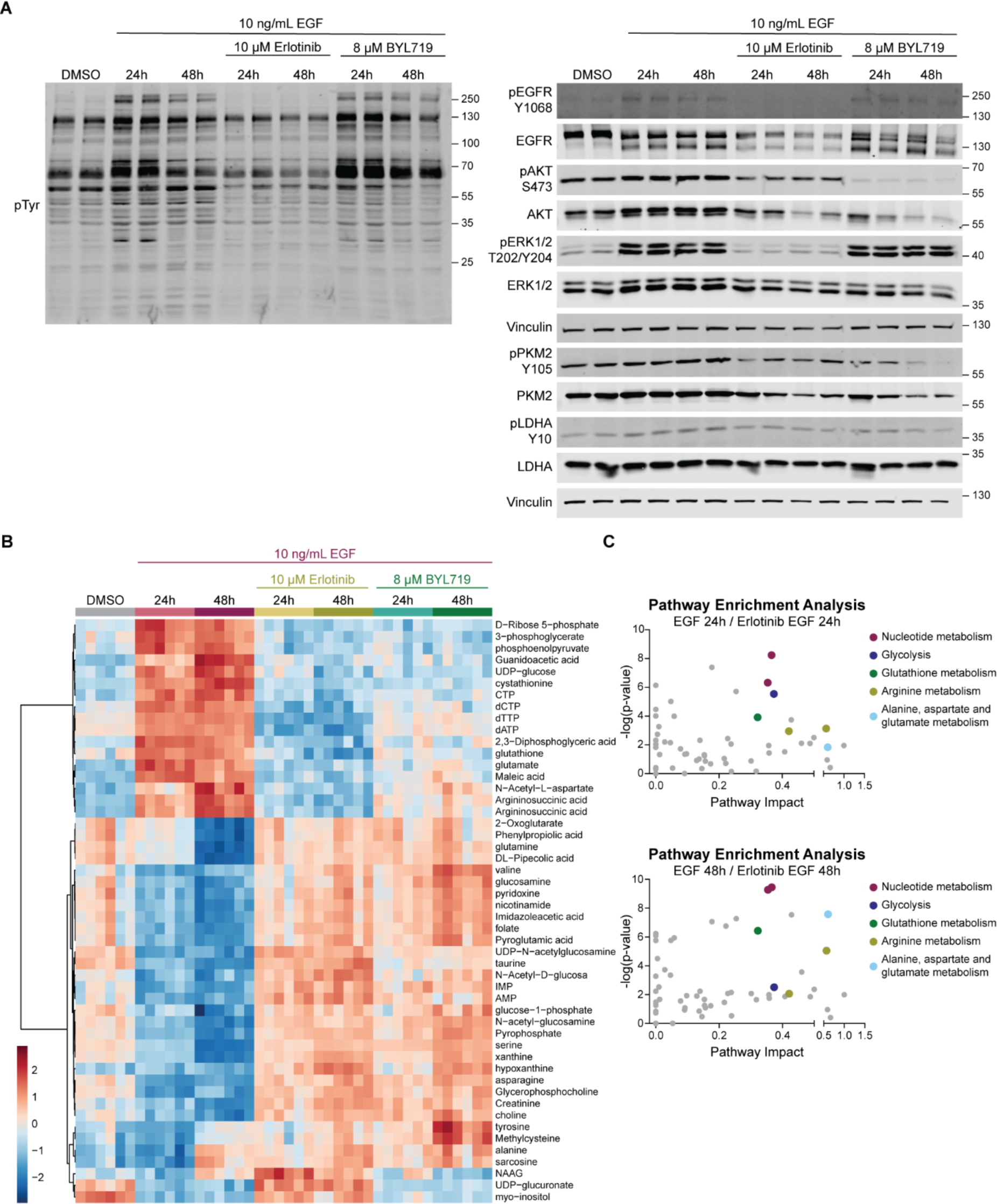
Metabolomics in MCF10A cells identifies metabolic changes downstream of activated EGFR. *A,* Immunoblots of total phosphotyrosine (pTyr), pEGFR^Y1068^, pAKT^S473^, pERK1/2^T202/Y204^, pPKM2^Y105^ and pLDHA^Y10^ in MCF10A cells after treatment with 10 ng/mL EGF for 5 minutes, 24 hours or 48 hours with or without a 10-minute pre-treatment with erlotinib (10 µM) or BYL719 (8 µM). Cells were seeded and harvested in parallel with the metabolomics experiment in *B*. *B,* Heatmap showing abundance of top 50 polar metabolites in MCF10A cells after treatment with 10 ng/mL EGF for 24 or 48 hours with or without a 10-minute pre-treatment with erlotinib (10 µM) or BYL719 (8 µM) compared to DMSO. Heatmap was generated using log10 transformation and Pareto scaling on MetaboAnalyst 5.0. *C,* Pathways enriched in metabolites altered after 24 or 48 hours of EGF stimulation compared to erlotinib pre-treatment. Plots were generated using the enrichment analysis feature of MetaboAnalyst 5.0.

### Integrated analysis of phosphoproteomic and metabolomic datasets identifies metabolic enzymes with functionally relevant pTyr sites

After performing independent analyses of the phosphoproteomic and metabolomic datasets, we evaluated overlap between these datasets by both manual analysis and by utilizing the Joint Pathway Analysis feature of MetaboAnalyst^27^. Metabolic enzymes and pathways of interest met the following criteria: (1) metabolic enzymes had EGF-responsive pTyr sites that were detected by our phosphoproteomic analysis, and (2) metabolism was altered in EGF-stimulated versus erlotinib-pre-treated conditions. Both analysis methods converged on three key metabolic pathways that met those criteria: glycolysis, glutamine metabolism and glutathione metabolism. Tyrosine phosphorylation of glycolytic enzymes has been previously characterized, and our integrated analysis identified multiple pTyr sites on PKM (Y83, Y370) and PGAM1 (Y26, Y92) that corresponded with changes in glycolytic metabolites (Fig. 4*A-B*). To visualize the potential contributions of these pTyr sites to cellular metabolism, we mapped glycolytic metabolites colored by a heatmap scale indicating their relative abundance in EGF-stimulated versus erlotinib-pre-treated conditions. Each metabolic enzyme was similarly colored by a heatmap scale of fold-change in phosphosite abundance detected over the mean across all samples by phosphoproteomic analysis. Although we performed steady-state metabolomics, creating a pathway map with metabolite abundance allows us to infer pathway activity or flux. As expected, EGF stimulation enhanced flux through glycolysis compared to EGFR inhibition (Fig. 4*C*). Identification of these previously characterized pTyr sites on glycolytic enzymes further validated this approach for identifying functionally relevant pTyr sites.

**Figure 4.**
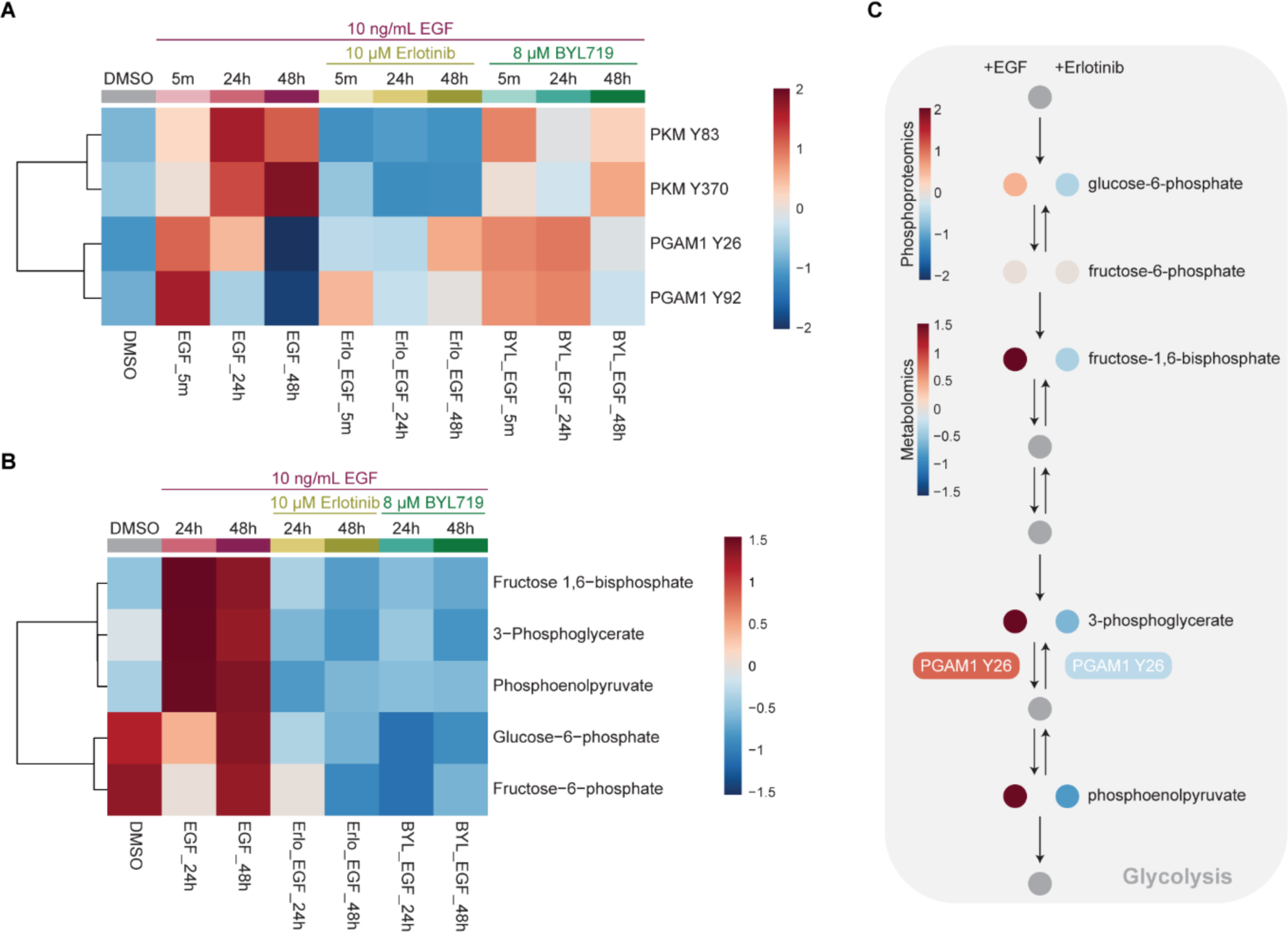
Integrated analysis of phosphoproteomic and metabolomic datasets identifies previously characterized pTyr sites on glycolytic enzymes. *A,* Heatmap showing fold-change in abundance of PKM Y83 and Y370 and PGAM1 Y26 and Y92 phosphosites detected over the mean across all samples in MCF10A cells after treatment with 10 ng/mL EGF for 5 minutes, 24 hours or 48 hours with or without a 10-minute pre-treatment with erlotinib (10 µM) or BYL719 (8 µM). *B,* Heatmap showing abundance of glycolytic metabolites in MCF10A cells after treatment with 10 ng/mL EGF for 24 or 48 hours with or without a 10-minute pre-treatment with erlotinib (10 µM) or BYL719 (8 µM) compared to DMSO. Heatmap was generated using log10 transformation and Pareto scaling on MetaboAnalyst 5.0. *C,* Schematic of glycolysis, highlighting the role of PGAM1. PGAM1 and polar metabolites detected by metabolomics are colored by a heatmap scale representing either fold-change in abundance detected over the mean from phosphoproteomic analyses at 5 minutes or fold-change in metabolite abundance compared to DMSO at 24 hours.

We also identified previously uncharacterized EGF-responsive pTyr sites on GLUD1 (Y512) and GSTP1 (Y64) that coincided with altered glutamine or glutathione metabolism, respectively (Fig. 5*A-C*). pTyr of these residues occurred as early as 5 minutes after EGF stimulation and for GSTP1, was sustained for 24 hours (Fig. 5*A*). Given the sustained phosphorylation of GSTP1, which matched the phosphorylation pattern of functionally significant pTyr sites on glycolytic enzymes, we aimed to validate this novel site. Mapping metabolites involved in glutathione metabolism showed that EGF stimulation resulted in an accumulation of glutamate and glutathione, which was reversed with EGFR inhibition (Fig. 5*D*). These data suggest that phosphorylation of GSTP1 Y64 is an inhibitory event that causes accumulation of glutathione and upstream intermediates. Together, this mapping approach integrates phosphoproteomic and metabolomic data to suggest potential roles of metabolic enzyme pTyr in cellular metabolism.

**Figure 5.**
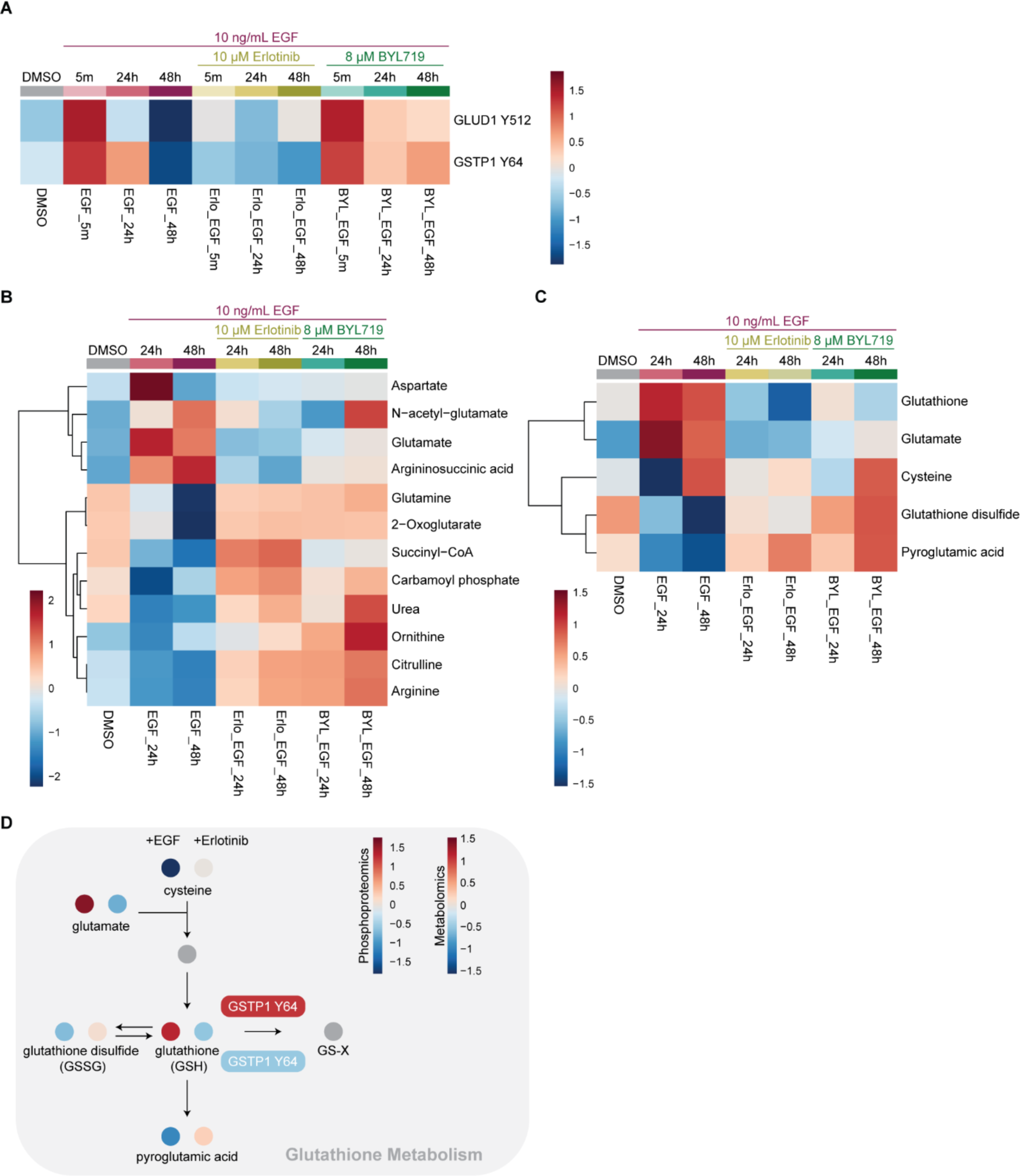
Integrated analysis of phosphoproteomic and metabolomic datasets identifies uncharacterized pTyr sites on GLUD1 and GSTP1. *A,* Heatmap showing fold-change in abundance of GLUD1 Y512 and GSTP1 Y64 phosphosites detected over the mean across all samples in MCF10A cells after treatment with 10 ng/mL EGF for 5 minutes, 24 hours or 48 hours with or without a 10-minute pre-treatment with erlotinib (10 µM) or BYL719 (8 µM). *B,* Heatmap showing abundance of metabolites related to GLUD1 in MCF10A cells after treatment with 10 ng/mL EGF for 24 or 48 hours with or without a 10-minute pre-treatment with erlotinib (10 µM) or BYL719 (8 µM) compared to DMSO. *C,* Heatmap showing abundances of metabolites related to GSTP1 (glutathione metabolism) in MCF10A cells after treatment with 10 ng/mL EGF for 24 or 48 hours with or without a 10-minute pre-treatment with erlotinib (10 µM) or BYL719 (8 µM) compared to DMSO. Heatmaps in *B* and *C* were generated using log10 transformation and Pareto scaling on MetaboAnalyst 5.0. *D,* Schematic of glutathione metabolism, highlighting the role of GSTP1. GSTP1 and polar metabolites detected by metabolomics are colored by a heatmap scale representing either fold-change in abundance detected over the mean from phosphoproteomic analyses at 5 minutes or fold-change in metabolite abundance compared to DMSO at 24 hours.

### Validation of hits from multi-omics analysis using a doxycycline-inducible CRISPR interference (CRISPRi) system

We next performed doxycycline-inducible CRISPR interference (CRISPRi) knockdown in MCF10A cells to deplete endogenous target metabolic enzymes with simultaneous inducible re-expression of mutant isoforms. This system allows for the interrogation of metabolic enzymes whose total knockout is not viable and prevents metabolic adaptions in response to metabolic enzyme depletion prior to re-expression of the enzyme. To characterize the roles of specific pTyr sites on metabolic enzyme activity, we overexpressed wild-type (WT), phosphomutant (tyrosine to phenylalanine, YtoF) or phosphomimetic (tyrosine to glutamate, YtoE) enzyme variants. We generated this system for PGAM1 Y26 as a positive control and for GSTP1 Y64. The efficiency of metabolic enzyme depletion in this system was variable, depending on the enzyme being targeted (Fig. 6*A*). However, PGAM1 and GSTP1 were both significantly overexpressed compared to endogenous enzyme expression levels (Fig. 6*B*).

**Figure 6.**
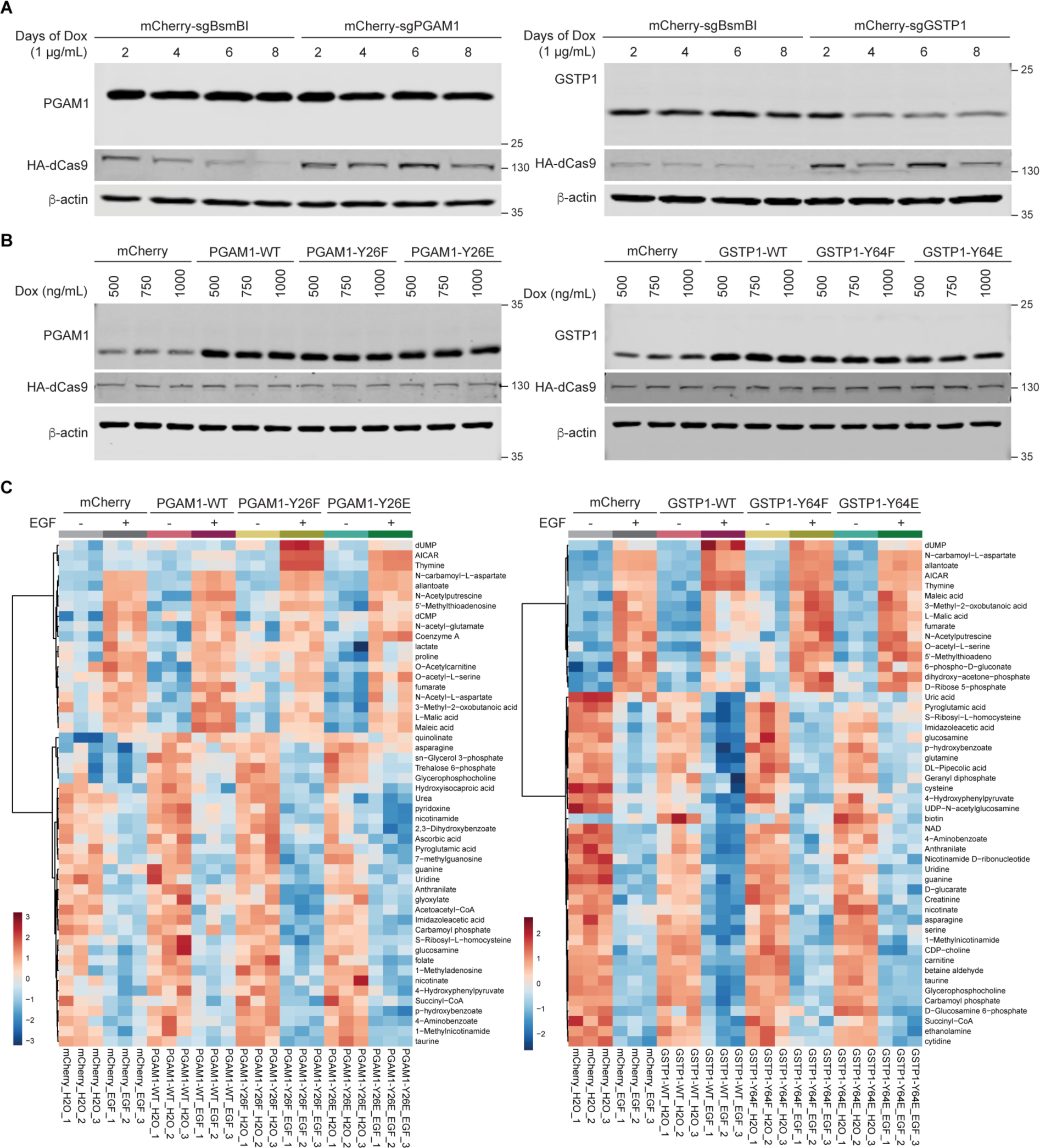
A doxycycline-inducible CRISPR interference (CRISPRi) system allows for functional validation of pTyr events on metabolic enzymes. *A,* MCF10A cells were transfected with Super PiggyBac transposase (SPT), HA-tagged KRAB-dCas9 and four pooled sgRNAs targeting the metabolic enzyme of interest or a control (sgBsmBI) and then transduced with lentivirus to overexpress mCherry. Immunoblots show metabolic enzyme (PGAM1, GSTP1), HA-dCas9 and β-actin expression levels after 2, 4, 6 or 8 days of treatment with 1 µg/mL doxycycline (dox). *B,* MCF10A cells were transfected as in *A* and then transduced with lentivirus for dox-inducible overexpression of mCherry or WT, YtoF or YtoE metabolic enzyme. Immunoblots show metabolic enzyme (PGAM1, GSTP1), HA-dCas9 and β-actin expression levels after 4 days of treatment with 500, 750 or 1000 ng/mL dox. *C,* Heatmap showing abundance of top 50 polar metabolites in MCF10A cells expressing CRISPRi constructs for PGAM1 or GSTP1 after pre-treatment with 500 ng/mL doxycycline for 3 days, followed by stimulation with 10 ng/mL EGF for 24 hours. Heatmap was generated using log10 transformation and Pareto scaling on MetaboAnalyst 5.0.

We performed steady-state polar metabolomics in MCF10A cells with inducible depletion of PGAM1 or GSTP1 and simultaneous re-expression of mCherry (control), WT, YtoF or YtoE PGAM1 or GSTP1 after 24 hours of treatment with EGF. Analysis of the top 50 altered metabolites for each metabolic enzyme showed the most significant differences between EGF stimulation and vehicle control. However, some metabolites were uniquely altered with overexpression of the phosphomutant (YtoF) or phosphomimetic (YtoE) compared to WT enzyme (Fig. 6*C*). To better visualize these changes, we focused on metabolites involved in glycolysis or glutathione metabolism for PGAM1 or GSTP1, respectively. Overexpression of phosphomutant (Y26F) PGAM1 impaired the EGF-induced increase in fructose-6-phosphate and glucose-6-phosphate observed in cells expressing WT PGAM1, whereas overexpression of phosphomimetic (Y26E) PGAM1 resulted in EGF-induced increases in all glycolytic metabolites (Fig. 7*A*). These changes were mapped onto the glycolytic pathway to highlight the effects of this phosphorylation site on flux through the pathway. As expected, since phosphorylation of PGAM1 Y26 has been shown to activate PGAM1, overexpression of the phosphomimetic (Y26E) enhanced glycolytic flux compared to WT or phosphomutant (Y26F) PGAM1 (Fig. 7*B*).

**Figure 7.**
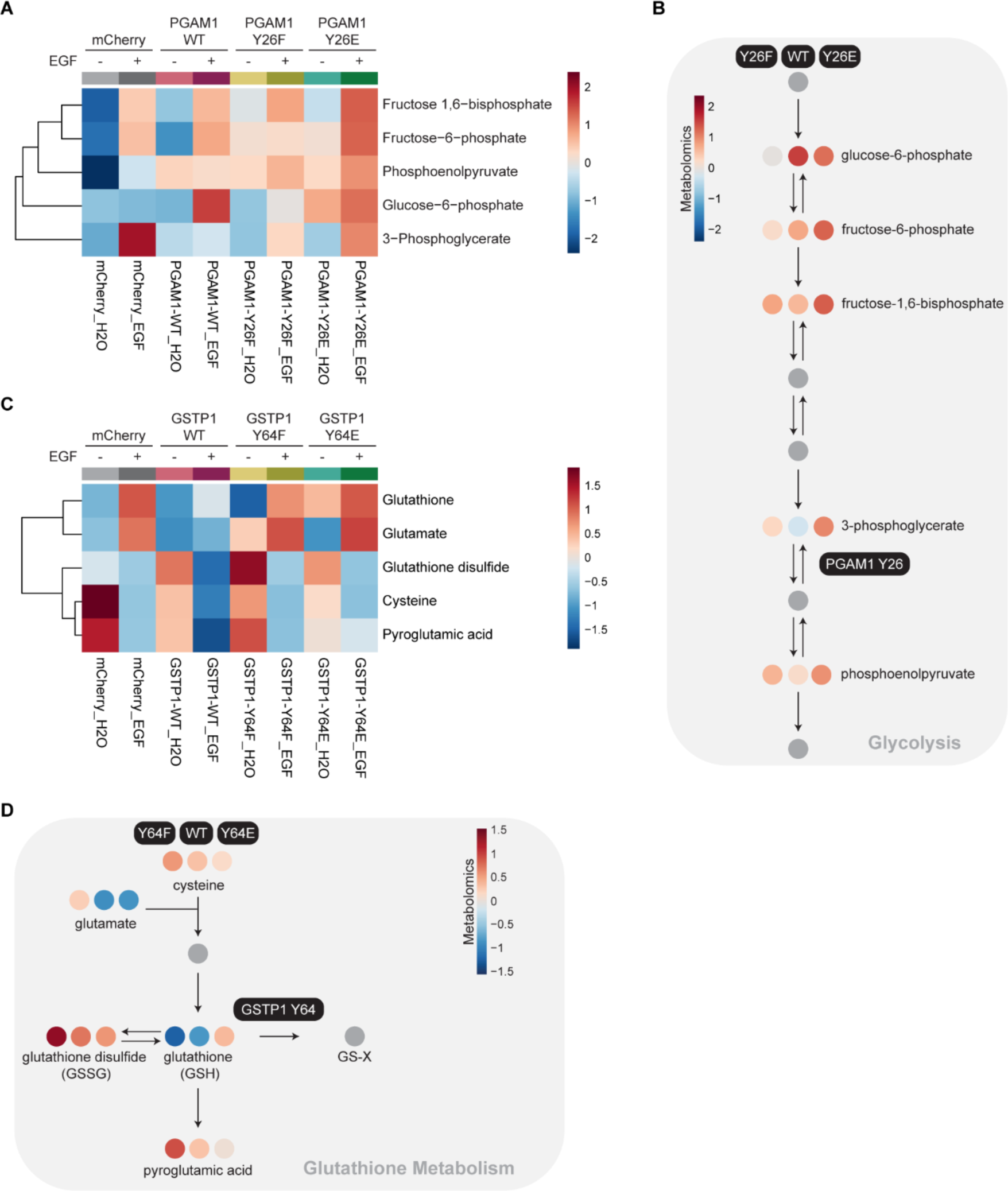
Metabolomics validates functional significance of pTyr sites on PGAM1 and GSTP1 identified by an integrated multi-omics analysis. *A,* Heatmap showing abundance of glycolytic metabolites in MCF10A cells expressing CRISPRi constructs for PGAM1 after pre-treatment with 500 ng/mL doxycycline for 3 days, followed by stimulation with 10 ng/mL EGF for 24 hours. Heatmap was generated using log10 transformation and Pareto scaling on MetaboAnalyst 5.0. *B,* Schematic of glycolysis, highlighting the role of wild-type (WT) versus phosphomutant (Y26F, Y26E) PGAM1. Polar metabolites detected by metabolomics are colored by a heatmap scale representing fold-change in metabolite abundance after EGF stimulation compared to the water (H_2_O) control. From left to right, circles represent metabolite abundance with expression of Y26F, WT or Y26E PGAM1. *C,* Heatmap showing abundance of metabolites involved in glutathione metabolism in MCF10A cells expressing CRISPRi constructs for GSTP1 after pre-treatment with 500 ng/mL doxycycline for 3 days, followed by stimulation with 10 ng/mL EGF for 24 hours. Heatmap was generated using log10 transformation and Pareto scaling on MetaboAnalyst 5.0. *D,* Schematic of glutathione metabolism, highlighting the role of wild-type (WT) versus phosphomutant (Y64F, Y64E) GSTP1. Polar metabolites detected by metabolomics are colored by a heatmap scale representing fold-change in metabolite abundance at baseline compared to the water (H_2_O) control. From left to right, circles represent metabolite abundance with expression of Y64F, WT or Y64E GSTP1.

We performed a similar analysis in GSTP1-expressing cells. Compared to WT, overexpression of phosphomutant (Y64F) GSTP1 decreased glutathione levels at baseline, while overexpression of phosphomimetic (Y64E) GSTP1 increased glutathione levels (Fig. 7*C*). Mapping these changes in glutathione levels suggests that phosphorylation of GSTP1 Y64 is inhibitory, since overexpression of phosphomimetic (Y64E) GSTP1 resulted in increased glutathione levels, even in the absence of EGF stimulation (Fig. 7*D*). These data highlight the potential for multi-omics approaches to rapidly identify functionally significant pTyr sites on metabolic enzymes.

### Computational prediction of kinases responsible for pTyr events

Since EGFR activates a cascade of downstream tyrosine kinases, sites identified from this multi-omics strategy are not necessarily direct substrates of EGFR. To identify potential kinases responsible for tyrosine phosphorylation of PGAM1 and GSTP1, we utilized a computational approach based on kinase substrate motifs to infer kinase activity in our phosphoproteomic dataset (Fig. 2). This approach utilizes knowledge of phosphorylated substrate proteins from our phosphoproteomic dataset to predict which kinases are likely to be active^28^. After EGF stimulation, EGFR and other human epidermal growth factor receptor (HER) family members, all of which contain very similar substrate motifs, were predicted to be active (Fig. 8*A*). Interestingly, other RTKs (RET, MET, FGFR2, EPHA3) and NRTKs (CSK, ACK, FES) were also predicted to be active after EGF stimulation, suggesting that PGAM1 and GSTP1 could be phosphorylated by these kinases downstream of EGFR (Fig. 8*A*). Similar analyses were performed in the erlotinib and BYL719 pre-treatment conditions. Erlotinib pre-treatment inhibited most EGF-activated kinases (Fig. 8*B*). BYL719 pre-treatment inhibited some kinases but retained the activity of others (Fig. 8*C*). In addition to predicting kinase activity, we also predicted EGFR substrates in our phosphoproteomic data. Putative EGFR substrates included autophosphorylation sites on EGFR and pTyr sites on other tyrosine kinases (Fig. 8*D*). Interestingly, PKM Y83 is a predicted EGFR substrate, suggesting that this approach can predict kinases responsible for phosphorylation events on metabolic enzymes (Fig. 8*D*). Altogether, we present a workflow for performing multi-omics analyses with joint pathway analysis to prioritize hits for downstream validation by multiple methods, including CRISPRi and computational prediction approaches (Fig. 8*E*).

**Figure 8.**
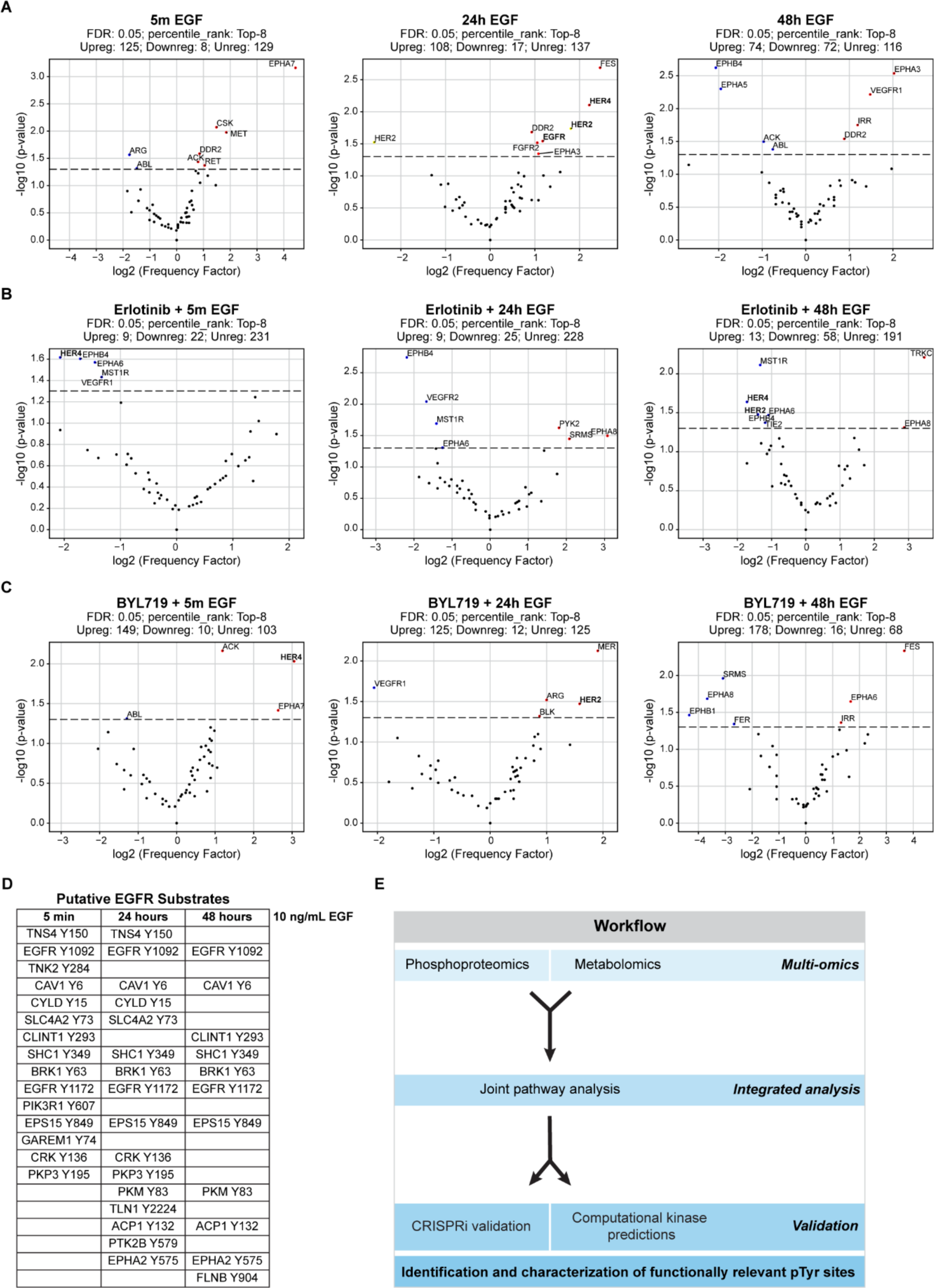
Prediction of putative kinases and substrates from phosphoproteomic data. *A-C,* Volcano plots of kinases predicted to be active after EGF stimulation (*A*), erlotinib treatment (*B*) or BYL719 treatment (*C*) based on the phosphoproteomic dataset in Fig. 2. The log2 frequency factor is plotted against the -log10 of the p-value. These enrichments were determined using one-sided exact Fisher’s tests. EGFR and HER family members with similar substrate motifs are highlighted in bold. *D,* Putative EGFR substrates from the phosphoproteomic data in Fig. 2 were predicted after 5 minutes, 24 hours or 48 hours of 10 ng/mL EGF stimulation. Putative substrates are listed in rank order for each time point. *E,* A model of the workflow developed: multi-omics platforms were utilized and analyzed jointly to identify targets, which were validated by downstream methods to rapidly identify pTyr sites on metabolic enzymes with functional significance.

## Discussion

Technological advances in proteomics, including the development of pTyr enrichment methods for phosphoproteomics, have enabled the study of formerly elusive tyrosine phosphorylation events. However, the tyrosine phosphoproteome remains largely understudied, and the contributions of specific pTyr events to cellular metabolism and disease states remain poorly understood. We developed a multi-omics strategy to simultaneously detect pTyr events (phosphoproteomics) and readout the functional effects of these pTyr events (metabolomics). We performed parallel phosphoproteomics and metabolomics in EGF-stimulated or EGFR-inhibited MCF10A cells and analyzed these datasets together to map the metabolic tyrosine phosphoproteome. Joint pathway analysis by MetaboAnalyst confirmed manual analysis of these datasets and highlighted key metabolic pathways with pTyr-regulated enzymes and altered metabolism in response to EGF stimulation versus EGFR inhibition, including glycolysis, glutamine metabolism and glutathione metabolism. Metabolic enzymes in these pathways are regulated by pTyr in response to EGF stimulation and include PKM, PGAM1, GLUD1 and GSTP1. By mapping the detected metabolites in these pathways in the context of EGF stimulation or EGFR inhibition, we inferred pathway flux and the functional consequences of these pTyr sites. For example, phosphorylation of PGAM1 Y26 has been shown to enhance substrate binding and increase enzymatic activity^9,16^. EGF stimulation resulted in phosphorylation of PGAM1 Y26 and coincided with increased phosphoenolpyruvate abundance, suggesting that phosphorylation of Y26 activates PGAM1 (Fig. 4*A-B*). These data confirm the findings of previous studies and further support a multi-omics approach for the functional characterization of metabolic enzyme pTyr sites.

In addition to identifying previously characterized pTyr sites on PKM and PGAM1, we also identified pTyr sites on GLUD1 and GSTP1 that have been detected in multiple HTP studies on PhosphoSitePlus but have not been extensively characterized. Both Y512 of GLUD1 and Y64 of GSTP1 were phosphorylated in response to EGF stimulation, and this was abolished with erlotinib pre-treatment or sustained with BYL719 pre-treatment. The phosphorylation pattern of GSTP1 Y64 mirrored that of the pTyr sites on PKM and PGAM1, supporting its functional significance. Phosphorylation of GSTP1 Y64 coincided with increased glutathione and glutamate levels (Fig. 5*A-B*). Overexpression of phosphomutant (Y64F) GSTP1 caused an accumulation of upstream metabolites, suggesting that phosphorylation of GSTP1 Y64 is inhibitory (Fig. 7*C-D*). These findings are consistent with previous reports suggesting that GSTP1 is phosphorylated downstream of EGFR^29,30^. Here, we provide further characterization of GSTP1, showing that pTyr of specifically Y64 likely inhibits GSTP1 and promotes glutathione accumulation.

Identifying and characterizing pTyr sites on metabolic enzymes is critically important for understanding both normal cell physiology and disease states. Phosphorylation of PKM2 Y105 inhibits formation of the active PKM2 tetramer, thereby inhibiting PKM2 activity. Loss of PKM2 activity promotes the Warburg effect, a phenomenon in which cancer cells favor metabolism by glycolysis, rather than oxidative phosphorylation^9,17,18^. While the benefits of the Warburg effect for cancer cell metabolism continue to be uncovered, the importance of pTyr of PKM2 to mediate this effect is undisputed^31^. We propose that characterizing the approximately 1000 other metabolic enzymes with pTyr sites will provide essential insights into cancer cell metabolism and highlight avenues for potential cancer therapies.

Targeting cancer metabolism has emerged as an attractive therapeutic strategy, since the altered metabolism of cancer cells might offer a therapeutic window over normal cells^32^. Furthermore, targeting metabolism downstream of RTK signaling may limit toxicities and resistance mechanisms. TKIs have been widely used in cancer treatment with variable success. Apart from mutant-specific inhibitors, dose-limiting toxicities of TKIs have limited their clinical utility^11^. Identifying and targeting metabolic enzymes downstream of oncogenic RTK signaling could prevent toxicities associated with inhibition of other RTK-mediated processes while also limiting acquired resistance. In addition, recent advances in drug development have identified methods of targeting aberrant pTyr. Phosphatase recruiting chimeras (PhoRCs) are heterobifunctional molecules that have been developed to target a phosphatase to a specific protein of interest^33^. While PhoRCs have not yet been developed for clinical use, their promise is supported by the preliminary success of another class of heterobifunctional molecules, proteolysis targeting chimeras (PROTACs). The mechanism of action of PhoRCs is especially attractive for targeting proteins with essential roles in normal cells, such as metabolic enzymes. By impairing their phosphorylation status, rather than degrading or inhibiting the metabolic enzyme, there is potential for therapeutic efficacy without toxicity. Targeting metabolic enzymes with aberrant pTyr with PhoRCs could selectively kill cancer cells, while sparing normal cells and limiting resistance and toxicities.

Given the abundance of open source, publicly-available omics datasets, methods to extract and prioritize biologically relevant targets are needed. Recent work has developed an atlas of substrate specificities for the human serine/threonine and tyrosine kinomes, enabling the computational prediction of kinases responsible for every serine, threonine or tyrosine phosphorylation event^28,34^. Integrating this resource with available datasets allows for rapid assignment of kinases to substrates and can identify novel biology from existing data. Furthermore, utilizing these atlases in parallel with functional experiments can rapidly characterize functionally significant targets. Yet, unlike many serine/threonine kinases, tyrosine kinases lack stringent substrate motifs, limiting computational predictions of kinase activity and highlighting the need for integration of these resources with experimental data. Here, we have described a multi-omics approach for the identification of pTyr sites on metabolic enzymes downstream of EGFR. This strategy could be extended to identify other RTK substrates or other post-translational modifications. Similarly, phosphoproteomics could be performed in parallel with other methods for functional readout of protein phosphorylation. These global approaches will facilitate mapping the dark phosphoproteome and could likely uncover new drug targets for diseases driven by aberrant protein phosphorylation.

## Methods

### Cell lines

The following commercially available cell lines were used: MCF10A (ATCC, CRL-10317) and HEK 293T (ATCC, CRL-11268). MCF10A cells were cultured in standard MCF10A growth medium without antibiotics (DMEM/F12 medium (Wisent Bioproducts, 319-075-CL), 5% horse serum (Gemini Bio, 100508), 10 µg/mL insulin (ThermoFisher Scientific/Gibco, A11382II), 0.5 mg/mL hydrocortisone (Sigma-Aldrich, H4001), 20 ng/mL EGF (R&D Systems, 236-EG-01M) and 100 ng/mL cholera toxin (List Biological Laboratories, 100B)). For phosphoproteomics and metabolomics, MCF10A cells were seeded and maintained in “assay media” without antibiotics (DMEM/F12 medium, 2% horse serum, 10 µg/mL insulin, 0.5 mg/mL hydrocortisone and 100 ng/mL cholera toxin). See the CRISPRi section below for information about the maintenance of MCF10A cells modified by CRISPRi.

### Phosphoproteomics

Cell seeding, drug treatments and protein harvest. Cells were plated at 175,000 cells/mL in tissue culture-treated 15-cm dishes in MCF10A “assay media,” which contains the standard components of MCF10A media with the following modifications: horse serum is lowered to 2% and no EGF is added. Four technical replicates were seeded for analysis by phosphoproteomics. The next day stocks of the indicated compounds were prepared in water (EGF) or DMSO (erlotinib and BYL719), and cells were treated for 5 minutes, 24 hours or 48 hours with or without a 10-minute pre-treatment with erlotinib or BYL719. Treatments were spiked into the media, and media was not changed over the 48-hour period. A master mix of each drug stock was prepared and used to treat all conditions across replicates, stored at −80°C between treatments. At the endpoint, cells were harvested rapidly by pouring the media off each plate and rinsing them with cold 1X PBS. Cells were scraped on wet ice in 1 mL of fresh 8M urea (Sigma-Aldrich, U5128) in milliQ water, and each sample was moved to a 15 mL conical tube.

### Protein digestion, cleanup and multiplexing

Disulfide bonds were reduced with 10 mM dithiothreitol (DTT; Sigma-Aldrich, D9779) at 56°C for 1 hour and alkylated with 55 mM iodoacetamide (IAA; Sigma-Aldrich, I1149) at room temperature protected from light for 1 hour. Samples were diluted 10-fold (v:v) with 100 mM ammonium acetate, pH 8.9, to bring down the urea concentration below 1 M. Protein was digested using sequencing-grade trypsin (Promega, V511) at 1:50 (20 µg trypsin:1 mg protein) overnight at room temperature. Digestion was quenched with 1:40 (v:v) of 100% glacial acetic acid (Sigma-Aldrich, 338826). Peptide desalting and cleanup was performed using SepPak Light C-18 cartridges. Peptide volume was reduced using a SpeedVac concentrator (ThermoFisher Scientific). Sample aliquots of 200 µg were lyophilized for multiplexing. Dried peptides were resuspended in 50 mM N-2-hydroxyethylpiperazine-N’-2-ethanesulfonic acid (HEPES), pH 8.5, multiplexed using TMT 11-plex at 1:2 (w:w) of samples:TMT and labeled for 1 hour at room temperature. Samples were then quenched with 5% hydroxylamine (ThermoFisher Scientific, 90115) for 15 minutes before combining all samples. Labeled peptides were dried in the SpeedVac concentrator and stored at −80°C until phosphoenrichment steps.

### Immunoprecipitation, phosphoenrichment and mass spectrometry

For phosphotyrosine (pTyr) immunoprecipitation (IP), protein G agarose beads were conjugated with 24 µg of 4G10 anti-pTyr and 6 µg of PT-66 anti-pTyr antibodies in IP buffer (100 mM Tris-HCl, 0.3% Nonidet P-40, pH 7.4) with 10 mM imidazole for 6 hours at 4°C. Dried, TMT-labeled peptides were resuspended in IP buffer and incubated with bead-conjugate overnight. The supernatants from the pTyr IPs were collected for crude protein analysis. Beads were washed with IP buffer, and peptides were eluted with 0.2% trifluoroacetic acid. Eluted peptides were further enriched using the High-Select Fe-NTA phosphopeptide enrichment kit (ThermoFisher Scientific, A32992). Enriched phosphopeptides were loaded onto an analytical column (ID = 50 µm) packed with 5 µM C-18 beads. Phosphopeptide separation was performed via liquid chromatography (pTyr IP: 200-minute gradient with 70% acetonitrile in 0.2 mol/L acetic acid at a flowrate of 0.2 mL/minute), and data-dependent acquisition of peptides was done on an Orbitrap Exploris 480 mass spectrometer with MS1 scan settings of: m/z range = 350-2000, resolution=60000, AGC target = 3e6, maximum injection time = 50ms. The most intense ions in 3s cycle time were isolated for fragmentation using higher energy collusion dissociation (HCD) with settings: resolution=60000, AGC target = 1e5, maxIT=250ms, isolation window = 0.4m/z, collisional energy = 33%, dynamic exclusion = 20s Crude supernatant from the IP was diluted 1:1000 (v:v) and separated with chromatography settings of: 140-minute gradient with 70% acetonitrile in 0.2 mol/L acetic acid at a flowrate of 0.2 mL/minute. Data-dependent acquisition of peptides was performed on a Q Exactive Plus mass spectrometer as described previously^35^.

### Phosphoproteomics data processing

Each biological replicate was run as its own multiplex with a bridge sample to correct for run-to-run variability. Runs were searched using Proteome Discoverer 3.0, and raw peptide-spectrum match (PSM) files were exported for analysis. Each PSM was filtered based on the following cutoffs: difference in ppm of identified mass is [-10-10], expectation value < 0.05 and ion score ≥ 15. PSMs were removed if they were not observed in at least 20% of samples or if their combined TMT intensity across all samples was < 1500. Missing values were imputed using the minimum intensity of the corresponding MS1 scan, such that any empty value was assumed to be at/below this minimum value and thus was not fragmented for MS/MS. For each peptide, TMT intensities of all matching PSMs were summed, and the mean across all samples was calculated. To account for sample-to-sample variability, the summed peptides from crude supernatant runs were divided by the row mean (mean center intensities). Then, the column mean was calculated for each TMT channel and used as a correction factor for variabilities in sample processing steps (0.7-1.2 is acceptable). After filtering the PSM files for a given run, each phosphosite was divided by the normalization value obtained from the supernatant sample from the corresponding TMT channel. Then, row means were calculated for each phosphosite across TMT channels to standardize each phosphosite. This standardized data was then used to perform hierarchical clustering and sample correlation analysis and was plotted as a heatmap.

### Phosphoproteomics data analysis

All data analysis was performed using Python and GraphPad PRISM 10. Python packages used include pandas, NumPy, SciPy, sklearn, GSEApy, matplotlib and seaborn. Hierarchical clustering was performed using Euclidean distance with complete metrics. Sample Spearman correlation and PCA analysis were performed using sklearn.

### Kinase motif-enrichment analysis of phosphoproteomics datasets

Kinase predictions were based on experimental biochemical data of their substrate motifs obtained for 78 tyrosine kinases. Their motifs were quantified into position-specific scoring matrices (PSSMs) and then applied computationally to score phosphorylation sites based on their surrounding amino acid sequences. These PSSMs were ranked against each site to identify the most favorable kinases. The phosphorylation sites detected in this study were scored by all the characterized kinase PSSMs, and their ranks were determined. For every non-duplicate, singly phosphorylated site, kinases that ranked within the top 8 out of the 78 total kinases were considered as biochemically predicted kinases for their respective phosphorylation site. For assessing kinase motif enrichment, we compared the percentage of phosphorylation sites for which each kinase was predicted among the downregulated/upregulated phosphorylation sites (sites with |log2 fold change| ≥1 and with FDR ≤0.05) with the percentage of biochemically favored phosphorylation sites for that kinase within the set of unregulated sites in this study (sites with |log2 fold change| <1 and with FDR >0.05). Statistical significance was determined using a one-sided Fisher’s exact test. Kinases that were enriched in both upregulated and downregulated analysis were excluded from downstream analysis.

## Polar metabolomics

### Cell seeding, drug treatments and metabolite harvest

Cells were plated at 175,000 cells/mL in tissue culture-treated 6-cm dishes in MCF10A “assay media,” which contains the standard components of MCF10A media with the following modifications: horse serum is lowered to 2% and no EGF is added. Three technical replicates were seeded for analysis by metabolomics and two for protein analysis by immunoblotting. This experiment was performed in biological duplicate and six samples per condition were included in the analysis. The next day stocks of the indicated compounds were prepared in water (EGF) or DMSO (erlotinib and BYL719), and cells were treated for 24 or 48 hours with or without a 10-minute pre-treatment with erlotinib or BYL719. Treatments were spiked into the media, and media was not changed over the 48-hour period. A master mix of each drug stock was prepared and used to treat all conditions across biological replicates, stored at −80°C between treatments. After 24 or 48 hours of treatment, plates were placed on wet ice and media was aspirated completely. Plates were washed once with 2 mL of cold PBS, aspirated completely and moved to dry ice. On dry ice, 1.5 mL of 80% MeOH stored at −80°C was added to each plate. Plates were scraped on wet ice, and 1.5 mL of each sample was moved to a 2 mL Eppendorf tube and vortexed for 30 seconds. Samples were centrifuged at 20,000 x g at 4°C for 5 minutes and then immediately dried under nitrogen gas. After drying was complete, samples were stored at −80°C. To prepare the samples for LC/MS analysis, dried samples were resuspended in 20 µL of mass spectrometry-grade water, centrifuged at 10,000 x g at 4°C for 3 minutes, and supernatants were moved to a mass spectrometry-compatible vial. 2 µL of each sample was pooled and used to generate a dilution curve for data analysis.

### Targeted LC/MS metabolomics

Samples (5-7 µL) were injected and analyzed using a hybrid 6500 QTRAP triple quadrupole mass spectrometer (AB/SCIEX) coupled to a Prominence UFLC HPLC system (Shimadzu) via selected reaction monitoring (SRM) of a total of 300 endogenous water-soluble metabolites for steady-state analyses of samples. Some metabolites were targeted in both positive and negative ion mode for a total of 311 SRM transitions using positive/negative ion polarity switching. ESI voltage was +4950V in positive ion mode and –4500V in negative ion mode. The dwell time was 3 ms per SRM transition and the total cycle time was 1.55 seconds. Approximately 9-12 data points were acquired per detected metabolite. Samples were delivered to the mass spectrometer via hydrophilic interaction chromatography (HILIC) using a 4.6 mm i.d. x 10 cm Amide XBridge column (Waters) at 400 µL/min. Gradients were run starting from 85% buffer B (HPLC grade acetonitrile) to 42% B from 0-5 minutes; 42% B to 0% B from 5-16 minutes; 0% B was held from 16-24 minutes; 0% B to 85% B from 24-25 minutes; 85% B was held for 7 minutes to re-equilibrate the column. Buffer A was comprised of 20 mM ammonium hydroxide/20 mM ammonium acetate (pH=9.0) in 95:5 water:acetonitrile. Peak areas from the total ion current for each metabolite SRM transition were integrated using MultiQuant v3.0.2 software (AB/SCIEX).

### Metabolomics data analysis

Metabolomics data was processed and analyzed in RStudio. Two metabolomics runs were analyzed separately, and data was combined for heatmap generation and pathway analyses. Metabolites were removed from the analysis if they were not detected in greater than 3 samples (the number of technical replicates). A correlation coefficient was calculated for each metabolite using the pooled sample dilution curve. Metabolites with a correlation coefficient less than 0.7 were removed from the analysis. For metabolomics of CRISPRi cells, the correlation coefficient cutoff was omitted. Metabolite abundances were then normalized to protein levels determined by Bio-Rad DC protein assay of protein samples harvested in parallel. Protein-normalized metabolite abundances were normalized to the DMSO control. Protein- and control-normalized metabolite abundances were used to generate heatmaps and perform pathway analyses. Heatmaps were generated using the statistical analysis feature of MetaboAnalyst 5.0 with a log10 transformation and Pareto scaling.

### Immunoblotting

Cells were washed once in cold 1X PBS (Boston BioProducts, BM-220) and collected on wet ice in 4°C RIPA lysis buffer (150 mM Tris-HCl, 150 mM NaCl, 0.5% (w/v) sodium deoxycholate (Sigma-Aldrich, D6750), 1% (v/v) NP-40 (Sigma-Aldrich, I3021), pH 7.5) containing 0.1% sodium dodecyl sulfate (SDS; AmericanBio, AB01920-00500) and 1X HALT protease and phosphatase inhibitor cocktail (Fisher Scientific, PI78443), added just before use. Plates were scraped into 1.5 mL microcentrifuge tubes, vortexed and incubated on wet ice for 15 minutes. Samples were then centrifuged at 14,000 x rpm for 10 minutes at 4°C. The supernatants were transferred to new tubes, and protein concentrations were measured by the Bio-Rad DC protein assay (Bio-Rad, Reagent A: 5000113, Reagent B: 500-0114).

Sample concentrations were normalized using 2X SDS sample buffer (62.5 mM Tris pH 6.8, 2% SDS, 10% glycerol (Fisher Scientific G33-500), bromophenol blue (Fisher Scientific, BP115-25)). Cell lysates were boiled at 95°C for 5 minutes and stored at −20°C. Cell lysates were run by SDS-PAGE on 10% acrylamide gels in 1X running buffer (Boston BioProducts, BP-150). Proteins were transferred to nitrocellulose membranes at 100 volts for 90 min in cold 1X transfer buffer (Cell Signaling Technology, 12539S). Membranes were blocked with 5% (w/v) bovine serum albumin (BSA; Gold Biotechnology, A-420-100) in tris-buffered saline (TBS; Boston BioProducts, BM-301), rocking for 1 hour at room temperature. Membranes were washed briefly in TBST (TBST; Boston Bioproducts, IBB-181X) and incubated in primary antibodies diluted in 5% (w/v) BSA in TBST with 0.01% (w/v) sodium azide (Fisher Scientific, S227I-25), rocking at 4°C overnight. Membranes were washed three times for 10 minutes in TBST, rocking at room temperature. Then, membranes were incubated for 1 hour rocking at room temperature with fluorophore-conjugated secondary antibodies (LI-COR Biosciences). Membranes were washed two times for 10 minutes each in TBST and one time for 10 min in TBS and imaged with the LI-COR Odyssey CLx Imaging System (LI-COR Biosciences).

**Table 1.** Antibodies for immunoblotting.

|  | Antibody | Manufacturer | Identifier | Host | Dilution |
| --- | --- | --- | --- | --- | --- |
| Primary antibody | pTyr | Cell Signaling Technology | 9411 | Mouse | 1:1000 |
|  | pEGFR Y1068 | Cell Signaling Technology | 3777 | Rabbit | 1:1000 |
|  | EGFR | Cell Signaling Technology | 4267 | Rabbit | 1:1000 |
|  | pAKT S473 | Cell Signaling Technology | 4060 | Rabbit | 1:1000 |
|  | AKT | Cell Signaling Technology | 4691 | Rabbit | 1:1000 |
|  | pERK1/2 T202/Y204 | Cell Signaling Technology | 4370 | Rabbit | 1:1000 |
|  | ERK | Cell Signaling Technology | 4695 | Rabbit | 1:1000 |
|  | pPKM2 Y105 | Cell Signaling Technology | 3827 | Rabbit | 1:500 |
|  | PKM2 | Cell Signaling Technology | 4053 | Rabbit | 1:500 |
|  | pLDHA Y10 | Cell Signaling Technology | 8176 | Rabbit | 1:500 |
|  | LDHA | Cell Signaling Technology | 3582 | Rabbit | 1:500 |
|  | PGAM1 | Cell Signaling Technology | 12098 | Rabbit | 1:500 |
|  | GSTP1 | Cell Signaling Technology | 3369 | Mouse | 1:500 |
|  | HA-Tag | Cell Signaling Technology | 2367 | Mouse | 1:1000 |
| | β-actin | Cell Signaling Technology | 4970 | Rabbit | 1:3000 |
|  | Vinculin | Cell Signaling Technology | 13901 | Rabbit | 1:1000 |
| Secondary antibody | Goat anti-Rabbit IgG (H + L) | LI-COR | 926-32211 | Goat | 1:20,000 |
|  | Goat anti-Mouse IgG (H + L) | LI-COR | 926-32210 | Goat | 1:20,000 |

## CRISPR interference (CRISPRi) system

### Lentivirus production

HEK 293T cells were seeded in tissue culture-treated 10-cm plates to reach 80% confluency the following day. 24 hours after seeding, HEK 293T cells were transfected with plasmid DNA for PGAM1 or GSTP1 as follows. 10 µg of plasmid DNA was combined with 600 µL of Opti-MEM, 10 µg of psPAX2 (Addgene, 12260) and 5 µg of pMDG.2 (Addgene, 12259). 48 µL of 1 mg/mL polyethylenamine (PEI; Polysciences Inc, 24765-1) was added to the DNA mixture. The mixture was inverted, spun down briefly and incubated at room temperature for 20 minutes. After incubation, the mixture was added dropwise to a 10-cm dish of HEK 293T cells, mixed slowly and incubated at 37°C overnight. Approximately 16 hours after transfection, media was changed to 30% FBS-containing media. Lentivirus was harvested 48 hours after the media change by passage through a 0.45 µm filter and snap frozen.

### CRISPRi transfections

The CRISPRi system utilizes three main components: (1) the Super PiggyBac transposase vector (SPT), which was cloned into a pUC19 backbone with an ampicillin selection marker, (2) doxycycline (dox)-inducible HA-KRAB-dCas9 (Addgene, 126030) and (3) an sgRNA vector (Addgene, 126028) with the neomycin selection marker changed to blasticidin. The sgRNA vector was linearized by digesting with BsmBI to create sticky ends. Then, gene-specific sequences for PGAM1 and GSPT1, flanked with sticky ends, were ligated into the vector to circularize it again. MCF10A cells were seeded in tissue culture-treated 6-cm plates to be 80% confluent in 24 hours (500,000 cells/mL). The following day, MCF10A cells were transfected with 8 µg of DNA in tissue culture-treated 6-cm plates at a ratio of 4 µg of SPT:2 µg of HA-KRAB-dCas9:2 µg sgRNA (0.5 µg of 4 individual sgRNAs per gene were combined to make a pool). Just prior to transfection, MCF10A cell media was changed to 5 mL of fresh MCF10A media. The following reagents were combined into a tube and gently mixed: 250 µL of Opti-MEM, 2 µg of plasmid DNA, 2 µg of HA-KRAB-dCas9, 8 µg of SPT and 16 µL of P3000 reagent (2 µL/µg of DNA; ThermoFisher Scientific, L3000015). In another tube, 250 µL of Opti-MEM was combined with 10.1 µL of lipofectamine 3000 (2.5 µL/mL of final volume of media; ThermoFisher Scientific, L3000015) and gently mixed. Both tubes were incubated at room temperature for 5 minutes. After incubation, the contents of the first tube were dripped into the second tube, mixed gently (tubes were not vortexed) and incubated at room temperature for 15 minutes. After incubation, the DNA/lipofectamine mixture was added dropwise to MCF10A cells. The plate was tipped back and forth to mix evenly. Cells were incubated with transfection reagents for 48 hours. After 48 hours, cells were reaching confluency, but were not passaged, since the transfection efficiency is low. The transfection media was removed and changed to antibiotic-containing MCF10A media: 200 µg/mL hygromycin (Gibco, 10-687-010) and 5 µg/mL blasticidin (InvivoGen, ant-bl-1). Cells were selected for 48 hours, until the untransfected control cells were completely dead. Then, media was changed to MCF10A media with maintenance concentrations of antibiotics: 100 µg/mL hygromycin and 2 µg/mL blasticidin. Cells were allowed to recover and were passaged and frozen down.

### Lentiviral transduction of transfected cell lines

MCF10A cells that were successfully transfected with the dox-inducible CRISPRi system (SPT, HA-KRAB-Cas9 and sgRNA) were transduced with lentivirus for dox-inducible overexpression of wild-type (WT), phosphomutant (tyrosine to phenylalanine, YtoF) or phosphomimetic (tyrosine to glutamate, YtoE) PGAM1 or GSTP1. MCF10A cells were seeded in antibiotic-free MCF10A media in tissue culture-treated 6-well plates to be 50-60% confluent in 24 hours (175,000 cells/mL). The following day, MCF10A cells were transduced with 1 mL of lentivirus for expression of mCherry (control), WT, YtoF or YtoE enzyme. Polybrene (Sigma-Aldrich, 107689) was added at a final concentration of 10 µg/mL, and cells were incubated at 37°C for 24 hours. After 24 hours, the media was changed to antibiotic-containing MCF10A media: 100 µg/mL hygromycin, 2 µg/mL blasticidin and 2 µg/mL puromycin. Cells were allowed to recover and were passaged and frozen down when appropriate. After selection was complete, cells were maintained in media with 100 µg/mL hygromycin, 2 µg/mL blasticidin and 1 µg/mL puromycin. To knockdown the target metabolic enzyme and overexpress the transduced construct, cells were treated with 500-1000 ng/mL doxycycline (dox; Sigma-Aldrich, D3447) for 2-8 days, with dox replenished every 24 to 48 hours.

### Limitations of the study

Phosphoproteomics was performed using data-dependent analysis which limits detection of low abundance pTyr on metabolic enzymes that may not be sampled due to high intensity sites on signaling proteins. Similarly, the polar metabolomics analysis did not capture all metabolites directly linked to enzymes with pTyr measurements. Furthermore, the polar metabolomics analyses captured steady-state metabolite levels, and thus cannot be used to ascertain pathway flux. In our CRISPRi experiments, we only compared phosphomutants to wild-type, since the knockdown efficiency in MCF10A cells was not complete.

## Acknowledgements

This work was supported by the following grants: Ludwig Center at Harvard (A.T.), CA253097 (A.T.), NSF Graduate Research Fellowship (A.L.H.), Harvard University Landry Cancer Biology Fellowship (A.L.H.), Burroughs Wellcome Fund PDEP Fellowship (T.Y.T.), U-01 Diversity Supplement (T.Y.T.), National Institute of Health grants P01-CA120964 (L.C.C.), R35-CA197588 (L.C.C.), P01-CA117969 (L.C.C.), Claudia Adams Barr Program for Cancer Research Award (J.L.J.). The mass spectrometry work was partially funded by NIH grants 5P01CA120964 (J.M.A.) and 5P30CA006516 (J.M.A.).

## Author contributions

Conceptualization: A.L.H., A.T.; methodology: A.L.H., T.T., G.E.P., F.M.W., A.T.; investigation: A.L.H., T.T., G.E.P, J.M.A., J.L.J., T.M.Y.; resources: T.T., F.M.W.; writing-original draft: A.L.H., A.T.; writing-review: A.L.H., T.T., G.E.P., J.M.A., J.L.J., T.M.Y., L.C.C., F.M.W., A.T.; writing-editing: A.L.H., T.T., A.T.; supervision: A.T.

## Data Availability

Source data and annotated analysis workflows are available on the following OSF project website: https://osf.io/3xmub/. Phosphoproteomic data are available via ProteomeXchange with identifier PXD052217.

